# Targeting YAP mechanosignaling to ameliorate stiffness-induced Schlemm’s canal cell pathobiology

**DOI:** 10.1101/2023.09.08.556840

**Authors:** Haiyan Li, Megan Kuhn, Ruth A. Kelly, Ayushi Singh, Kavipriya Kovai Palanivel, Izzy Salama, Michael L. De Ieso, W. Daniel Stamer, Preethi S. Ganapathy, Samuel Herberg

## Abstract

Pathologic alterations in the biomechanical properties of the Schlemm’s canal (SC) inner wall endothelium and its immediate vicinity are strongly associated with ocular hypertension in glaucoma due to decreased outflow facility. Specifically, the underlying trabecular meshwork is substantially stiffer in glaucomatous eyes compared to that from normal eyes. This raises the possibility of a critical involvement of mechanotransduction processes in driving SC cell dysfunction. Yes-associated protein (YAP) has emerged as a key contributor to glaucoma pathogenesis. However, the molecular underpinnings of SC cell YAP mechanosignaling in response to glaucomatous extracellular matrix (ECM) stiffening are not well understood. Using a novel biopolymer hydrogel that facilitates dynamic and reversible stiffness tuning, we investigated how ECM stiffening modulates YAP activity in primary human SC cells, and whether disruption of YAP mechanosignaling attenuates SC cell pathobiology and increases *ex vivo* outflow facility. We demonstrated that ECM stiffening drives pathologic YAP activation and cytoskeletal reorganization in SC cells, which was fully reversible by matrix softening in a distinct time-dependent manner. Furthermore, we showed that pharmacologic or genetic disruption of YAP mechanosignaling abrogates stiffness-induced SC cell dysfunction involving altered cytoskeletal and ECM remodeling. Lastly, we found that perfusion of the clinically-used, small molecule YAP inhibitor verteporfin (without light activation) increases *ex vivo* outflow facility in normal mouse eyes. Collectively, our data provide new evidence for a pathologic role of aberrant YAP mechanosignaling in SC cell dysfunction and suggest that YAP inhibition has therapeutic value for treating ocular hypertension in glaucoma.

## Introduction

The Schlemm’s canal (SC) inner wall endothelium is supported by a discontinuous basal lamina and numerous cell-cell contacts with the trabecular meshwork (TM) (1). Jointly, the two tissues form the central functional unit of the conventional outflow tract (2, 3). Pathological cellular and extracellular alterations in this region are responsible for the characteristic increased outflow resistance in primary open-angle glaucoma, the leading cause of irreversible blindness (4, 5). Mounting evidence suggest that the TM from glaucoma eyes is stiffer compared to that from age-matched normal eyes (6-9). Consequently, SC cells in the diseased outflow tract are subject to increased biomechanical stress from the stiffened TM underneath. A pivotal study from Overby *et al.* (10) showed that SC cells *in vitro* alter their transcriptional profile and stiffen in response to increased substrate stiffness, with SC cells isolated from glaucomatous eyes exhibiting a strikingly enhanced stiffening response. Moreover, Vahabikashi *et al.* (9) reported higher *in situ* SC cell stiffness in glaucomatous eyes, concomitant with increased outflow resistance. Together, these data suggest that dysregulation of cellular mechanotransduction plays a central role in SC cell pathobiology in glaucoma. However, the detailed mechanisms and pathways involved remain largely unaddressed.

The transcriptional coactivator Yes-associated protein (YAP) (11) is a master regulator of cellular mechanotransduction (12, 13). YAP activity is modulated by an ever-expanding network of input cues (14). For instance, increased extracellular matrix (ECM) stiffness strongly promotes YAP nuclear localization, where it binds primarily to TEAD transcription factors (as YAP lacks DNA-binding domains) to drive stiff-responsive gene expression (15, 16). On one hand, YAP tightly regulates essential tissue functions via this mechanism; on the other hand, impairment of this process is strongly associated with various diseases (17). In the eye, YAP was found to be activated by glaucoma-associated stressors in corneal fibroblasts (18), as well as TM cells (19-28) and lamina cribrosa cells (29). A recent genome-wide association study further identified *YAP1* among previously unknown open-angle glaucoma-risk loci (30), suggesting a potential causal association with outflow dysfunction. To examine this specifically, we recently linked glaucoma-like SC cell dysfunction directly with aberrant YAP activity using transforming growth factor beta2 as disease stimulus (31). However, the role of YAP mechanosignaling in SC cell pathobiology in response to ECM stiffening remains poorly understood.

Using a modified version of our 3D ECM hydrogel (32) that facilitates on-demand and reversible stiffness tuning, we investigated how ECM stiffening modulates YAP activity in primary human SC cells, and whether targeted interference with YAP mechanosignaling attenuates SC cell pathobiology and increases *ex vivo* outflow facility in mouse eyes.

## Materials and Methods

### SC cell isolation and culture

Experiments using human donor eye tissue were approved by the SUNY Upstate Medical University Institutional Review Board (protocol #1211036) and performed in accordance with the tenets of the Declaration of Helsinki for the use of human tissue. Primary human SC cells were isolated from ostensibly normal donor corneal rims discarded after transplant surgery, as recently described (31), and cultured according to established protocols (33). Four normal, previously characterized (31) SC cell strains (HSC01, HSC02, HSC03, HSC09) were used in the present study (**Table. 1**). All SC cell strains were characterized based upon their typical spindle-like elongated cell morphology, expression of vascular endothelial-cadherin and fibulin-2, and lack of myocilin induction following exposure to dexamethasone (31). Different combinations of SC cell strains were used per experiment depending on cell availability, and all studies were conducted between cell passage 3-6. SC cells were cultured in low-glucose Dulbecco’s Modified Eagle’s Medium (DMEM; Gibco; Thermo Fisher Scientific) containing 10% fetal bovine serum (FBS; Atlanta Biologicals, Flowery Branch, GA, USA) and 1% penicillin/streptomycin/glutamine (PSG; Gibco) and maintained at 37°C in a humidified atmosphere with 5% CO_2_. Fresh media was supplied every 2-3 days.

**Table 1.** SC cell strain information.

| **ID** | **Sex** | **Age** | **Used in** |
| --- | --- | --- | --- |
| HSC01 | Male | 33 | Figs. 1, 2, 3, 4; Suppl. Figs. 1, 2, 3, 4, 5 |
| HSC02 | Male | 46 | Figs. 1, 2, 3; Suppl. Figs. 1, 2, 3, 4 |
| HSC03 | Female | 46 | Figs. 1, 2, 4; Suppl. Figs. 1, 2, 4, 5 |
| HSC09 | Female | 69 | Figs. 1, 3 |

### Hydrogel precursor solutions

Methacrylate-conjugated bovine collagen type I (Advanced BioMatrix, Carlsbad, CA, USA) was reconstituted in sterile 20 mM acetic acid to achieve 6 mg/ml. Immediately prior to use, 1 ml of the collagen solution was neutralized with 85 µl neutralization buffer (Advanced BioMatrix) according to the manufacturer’s instructions. Thiol-conjugated hyaluronic acid (Glycosil®; Advanced BioMatrix) was reconstituted in sterile diH_2_O containing 0.5% (w/v) photoinitiator (4- (2-hydroxyethoxy) phenyl-(2-propyl) ketone; Irgacure® 2959; Sigma-Aldrich, St. Louis, MO, USA) to achieve 10 mg/ml according to the manufacturer’s protocol. In-house expressed elastin-like polypeptide (thiol via K<u>C</u>TS flanks (32)) was reconstituted in chilled Dulbecco’s Phosphate-Buffered Saline (DPBS) to achieve 10 mg/ml and sterilized using a 0.2 µm syringe filter, or UV sterilized in the biosafety cabinet for 20 min before reconstituting in chilled DPBS to achieve 50 mg/ml. Sodium alginate (Sigma-Aldrich) was reconstituted in diH_2_0 to achieve 20 mg/ml and sterilized using a 0.2 µm syringe filter.

### Hydrogel preparation

Hydrogel precursors methacrylate-conjugated bovine collagen type I (3.6 mg/ml), thiol-conjugated hyaluronic acid (0.5 mg/ml with 0.025% (w/v) photoinitiator), and elastin-like polypeptide (2.5 mg/ml) - with or without additional sodium alginate (4 mg/ml) - were thoroughly mixed in an amber color tube on ice. Thirty microliters of the hydrogel solution were pipetted onto Surfasil-coated (Fisher Scientific) 18 × 18-mm square glass coverslips followed by placing 12-mm round glass coverslips on top to facilitate even spreading of the polymer solution. Hydrogels were crosslinked by exposure to UV light (OmniCure S1500 UV Spot Curing System; Excelitas Technologies, Mississauga, Ontario, Canada) at 320-500 nm, 2.2 W/cm^2^ for 5 s, according to our established protocols (27, 28, 31, 32, 34, 35). The hydrogel-adhered coverslips were removed with fine-tipped tweezers and placed hydrogel-side facing up in 24-well culture plates (Corning; Thermo Fisher Scientific).

### Hydrogel stiffening and softening

Hydrogel stiffening was achieved by incubating pre-formed ECM-alginate hydrogels in 100 mM CaCl_2_ (Thermo Fisher Scientific) in diH_2_O to crosslink alginate at 37°C for 1 h. Soft control ECM hydrogels without alginate were treated in the same manner (i.e., no alginate, no change in stiffness). *In situ* softening of Ca^2+^-stiffened ECM-alginate hydrogels was achieved by digestion with 50 µg/ml alginate lyase (Sigma-Aldrich) in serum-free DMEM with 1% PSG at 37°C for 1 h.

### SC cell seeding and treatments

SC cells were seeded at 2 × 10^4^ cells/cm^2^ on premade soft ECM hydrogels or Ca^2+^-stiffened ECM-alginate hydrogels and cultured in DMEM with 10% FBS and 1% PSG for 1-2 days until ∼80-90%. Then, SC cell-seeded hydrogels were cultured in serum-free DMEM with 1% PSG for 6 days subjected to the following treatments: 1) soft, 2) stiff, or 3) softened (i.e., stiff for 3 days, soft for 3 hours, 1 day, 3 days, or 5 days).

For pharmacological YAP inhibition experiments, SC cells on Ca^2+^-stiffened ECM-alginate hydrogels were cultured in serum-free DMEM with 1% PSG for 3 days subjected to the following treatments: 1) control (vehicle: 0.1% dimethyl sulfoxide (DMSO); Fisher Scientific) or 2) verteporfin (0.5 μM; Sigma-Aldrich).

For siRNA-mediated YAP (and TAZ) depletion experiments, SC cells were seeded at 2 × 10^4^ cells/cm^2^ on Ca^2+^-stiffened ECM-alginate hydrogels in DMEM with 10% FBS and 1% PSG. The following day, the cell culture medium was changed to antibiotic-free and serum-free DMEM and the samples were kept in culture for 24 h followed by transfection. Transfection was performed using a final concentration 3% (v/v) lipofectamine RNAimax (Invitrogen; Thermo Fisher Scientific) with 150 nM RNAi duplexes (custom oligonucleotides; Horizon/Dharmacon, Lafayette, CO, USA), according to the manufacturer’s instructions. Transfected SC cells were analyzed 2 d after transfection. ON-TARGET plus nontargeting siRNA were obtained from Dharmacon. Custom siRNAs were identical to those used in our recent study (27), based on validated sequences (36): YAP, sense, 5’-GACAUCUUCUGGUCAGAGA-3’, and YAP, anti-sense, 5’-UCUCUGACCAGAAGAUGUC-3’; TAZ, sense, 5’-ACGUUGACUUAGGAACUUU-3’, and TAZ, anti-sense, 5’-AAAGUUCCUAAGUCAACGU-3’.

### Hydrogel rheology analysis

Fifty microliters of hydrogel precursor solution were pipetted into custom 8 × 1-mm polydimethylsiloxane molds. All hydrogels were photocrosslinked and exposed to CaCl_2_; samples in the softened group were subsequently exposed to alginate lyase, as described above. A Kinexus rheometer (Malvern Panalytical, Westborough, MA, USA) fitted with an 8-mm diameter parallel plate was used to measure hydrogel viscoelasticity. To ensure standard conditions across all experiments, the geometry was lowered into the hydrogels until a calibration normal force of 0.02 N was achieved. Subsequently, an oscillatory shear-strain sweep test (0.1-60%, 1.0 Hz, 25°C) was applied to determine storage modulus (G’) and loss modulus (G”) in the linear region from N = 3 experimental replicates per group. Elastic modulus was calculated with E = 2 * (1 + v) * G′, where a Poisson’s ratio (v) of 0.5 for the ECM hydrogels was assumed (37).

### Immunocytochemistry analysis

SC cells atop hydrogels were fixed with 4% paraformaldehyde (PFA; Thermo Fisher Scientific) at room temperature for 20 min, permeabilized with 0.5% Triton™ X-100 (Thermo Fisher Scientific), blocked with blocking buffer (BioGeneX, Fremont, CA, USA), and incubated with primary antibodies followed by incubation with fluorescent secondary antibodies (**Table 2**); nuclei were counterstained with 4′,6′-diamidino-2-phenylindole (DAPI; Abcam, Waltham, MA, USA). Similarly, cells were stained with Phalloidin-iFluor 488 (Invitrogen; Thermo Fisher Scientific) or 594 (Abcam)/DAPI according to the manufacturer’s instructions. Coverslips were mounted with ProLong™ Gold Antifade (Invitrogen; Thermo Fisher Scientific) on Superfrost™ microscope slides (Fisher Scientific), and fluorescent images were acquired with an Eclipse N*i* microscope (Nikon Instruments, Melville, NY, USA).

**Table 2.**
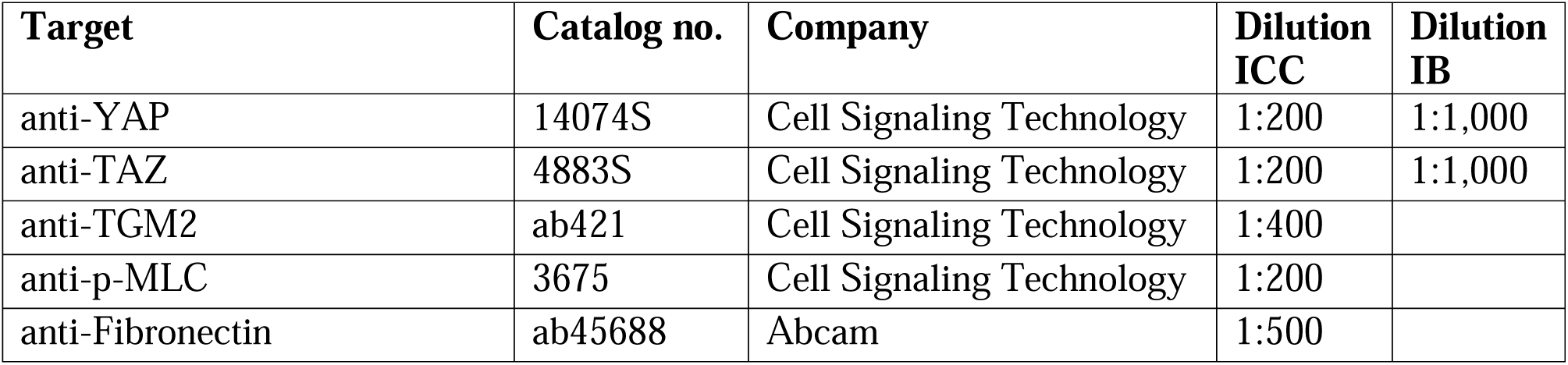
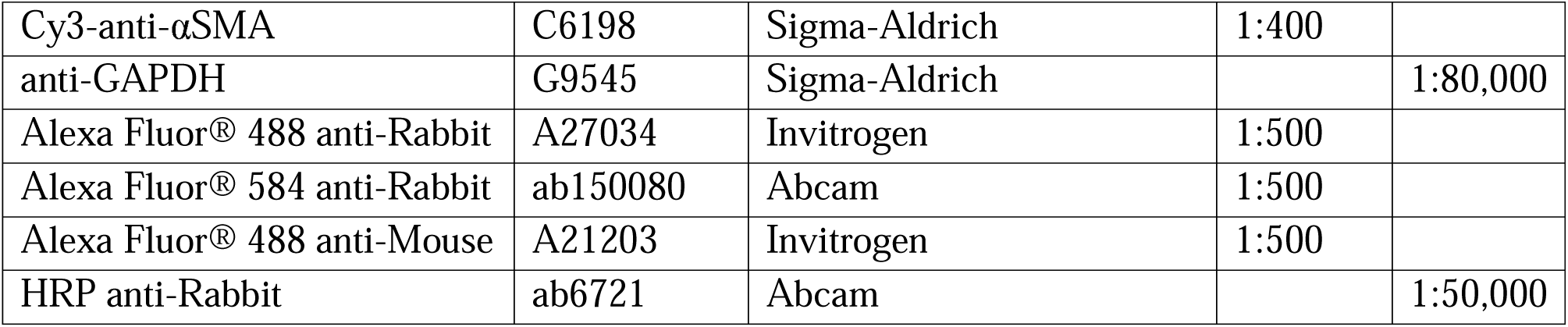
Antibody information.

All fluorescent image analyses were performed using FIJI software (38) (National Institutes of Health (NIH), Bethesda, MD, USA). The cytoplasmic YAP/TAZ intensity was measured by subtracting the overlapping nuclear (DAPI) intensity from the total YAP/TAZ intensity. The nuclear YAP/TAZ intensity was recorded as the proportion of total YAP/TAZ intensity that overlapped with the nucleus (DAPI). YAP/TAZ nuclear/cytoplasmic (N/C) ratio was calculated as follows: N/C ratio = (nuclear YAP/TAZ signal/area of nucleus)/(cytoplasmic signal/area of cytoplasm). Fluorescence intensity of F-actin, fibronectin (FN), transglutaminase 2 (TGM2), and phospho-myosin light chain (p-MLC) was measured with image background subtraction, followed by calculation of fold-change vs. control. Given the lack of defined alpha smooth muscle actin (αSMA) fibers in soft controls, we quantified the number of cells exhibiting αSMA signal/fibers (i.e., yes/no decision) per image after standardized background correction and calculated percent of αSMA-positive cells, as described recently (31). For each experiment, care was taken to include an equivalent number of cells across groups.

### Immunoblot analysis

Following siRNA transfection, protein was extracted from SC cells using lysis buffer (CelLytic^TM^ M, Sigma-Aldrich) supplemented with Halt™ protease/phosphatase inhibitor cocktail (Thermo Fisher Scientific). Equal protein amounts (10 µg), determined by standard bicinchoninic acid assay (Pierce; Thermo Fisher Scientific), in 4× loading buffer (Invitrogen; Thermo Fisher Scientific) with 5% beta-mercaptoethanol (Fisher Scientific) were boiled for 5 min and subjected to SDS-PAGE using NuPAGE™ 4-12% Bis-Tris Gels (Invitrogen; Thermo Fisher Scientific) at 120V for 80 min and transferred to 0.45 µm PVDF membranes (Sigma; Thermo Fisher Scientific). Membranes were blocked with 5% bovine serum albumin (Thermo Fisher Scientific) in tris-buffered saline with 0.2% Tween®20 (Thermo Fisher Scientific) and probed with primary antibodies followed by incubation with HRP-conjugated secondary antibodies (**Table 2**). Bound antibodies were visualized with the enhanced chemiluminescent detection system (Pierce) on autoradiography film (Thermo Fisher Scientific). Densitometry was performed using FIJI software (NIH) (38); data were normalized to GAPDH followed by calculation of relative change vs. control.

### Quantitative reverse transcription-polymerase chain reaction (qRT-PCR) analysis

Total RNA was extracted from SC cells atop hydrogels using PureLink RNA Mini Kit (Invitrogen; Thermo Fisher Scientific). RNA concentration was determined with a NanoDrop spectrophotometer (Thermo Fisher Scientific). RNA was reverse transcribed using iScript™ cDNA Synthesis Kit (BioRad, Hercules, CA, USA). One hundred nanograms of cDNA were amplified in duplicates in each 40-cycle reaction using a CFX 384 Real Time PCR System (BioRad) with annealing temperature set at 60°C, Power SYBR™ Green PCR Master Mix (Thermo Fisher Scientific), and custom-designed qRT-PCR primers (**Table 3**). Transcript levels were normalized to GAPDH, and mRNA fold-change calculated relative to mean control values using the comparative C_T_ method (39).

**Table 3.**
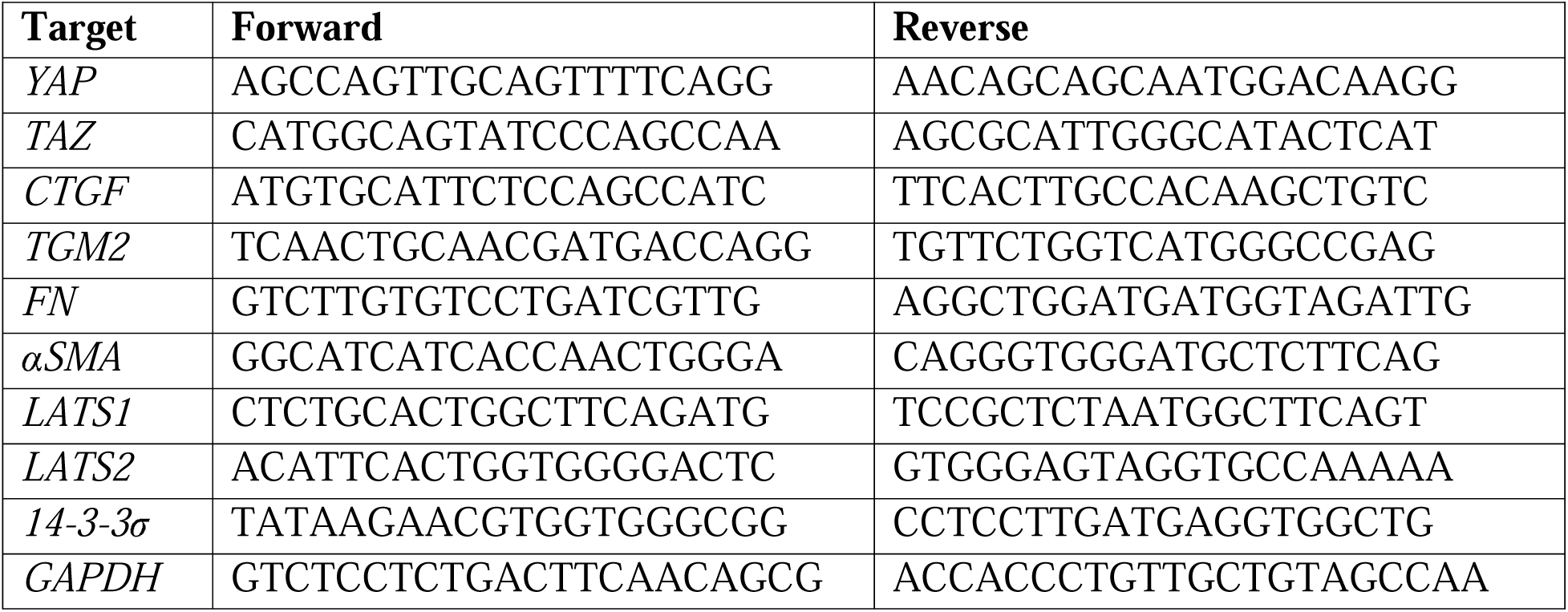
Oligonucleotide primer sequences (5’-3’) for qRT-PCR.

### Outflow facility analysis

Outflow facility was measured by *iPerfusion* (40) according to the institutional guidelines for the humane treatment of animals (Duke University IACUC protocol A128-21-06) and to the ARVO Statement for the Use of Animals in Ophthalmic and Vision Research, as described previously (40, 41). Nine C57BL/6J mice (both male and female, 3-6 months-old; Jackson Laboratory, Bar Harbor, ME, USA) were used for the study; mice were euthanized using isoflurane, and eyes were carefully enucleated and mounted on a stabilization platform located in the center of a perfusion chamber using a small amount of cyanoacrylate glue (Loctite, Westlake, OH, USA). The temperature-controlled perfusion chamber (35°C) was filled with prewarmed DPBS with added 5.5 mM D-glucose (DBG), submerging the eyes. To facilitate drug exchange (42, 43), two microneedles per eye were used: one needle connected to the syringe pump and used for the drug exchange and the other needle for the perfusion. The exchange needles were filled with DPBS and the perfusion needles were filled with filtered DBG containing verteporfin (10 µM) or vehicle control (DMSO). Of note, drug and control solutions were masked prior to shipment to Duke University and alternated between eyes of different mice (i.e., equal numbers of OD and OS eyes were treated with drug and control). The perfusion needles were connected to the perfusion system. Each pair of needles were held together with a plastic spacer and mounted on a micromanipulator to facilitate a dual needle cannulation, guided by a stereomicroscope. Following cannulation, eyes were acclimatized at 8 mmHg for 20 minutes. The syringe pump was then activated and fluid was withdrawn (via exchange needle) from the anterior chamber at a rate of 5 µl/min for 20 min. As fluid was withdrawn via the exchange needle, the perfusion needles delivered filtered DBG with and without verteporfin or vehicle control into the anterior chambers. After the exchange, the eyes underwent another 30-minute acclimation at 8 mmHg before starting facility measurement, which consisted of 9 steps that start at 5 mmHg, increase 1.5 mmHg each step until reaching 17 mmHg, then dropping down to 8 mmHg for the last step. Stable flow rate (Q) and pressure (P) averaged over 4 min at each pressure step were used for data analysis. A nonlinear flow-pressure model [Q = C_r_(P/P_r_)^β^P] that accounts for the pressure dependence of outflow facility in mice (β) was fit to the flow-pressure data using nonlinear regression, yielding the facility C_r_ evaluated at P_r_ = 8 mmHg, a pressure that approximates the physiological pressure drop across the conventional outflow pathway in living mice.

### Histology and immunohistochemistry analysis

Following *ex vivo* perfusion, eyes were fixed with 4% PFA overnight at 4°C. PFA was then removed, and eyes were washed twice with DPBS. Three pairs of eyes (both control and verteporfin-treated) showing good outflow facility response to the drug (i.e., > 40%) were used for histology and immunohistochemistry studies. Eyes were hemisected along the equator and the posterior segments and lenses removed. The anterior segments were then cut into four quadrants.

For histology studies, two quadrants were used. Each quadrant was embedded in Epon resin and 0.5 µm semi-thin sections were cut, stained with 1% of methylene blue and examined by light microscopy (Axioplan2, Carl Zeiss MicroImaging, Thornwood, NY, USA). To identify gross morphological differences between control and treated eyes, a trained masked observer (R.A.K) measured both the circumferential size and length of each SC lumen and counted the number of giant vacuoles per SC in each sample. All analyses were performed using ImageJ software (NIH). The circumferential size of SC was measured using the freehand line drawing tool. The length of SC lumen was measured using the straight-line drawing tool and the number of giant vacuoles present along the inner wall of SC were counted manually.

For immunohistochemistry studies, the remaining two quadrants were used. Each quadrant was placed in 30% sucrose overnight at 4°C, embedded in Tissue-Plus^TM^ O.C.T. Compound (Fisher Scientific) in a 10 × 10 × 5-mm cryomold, and placed in the -80°C freezer. Blocks were equilibrated at -20°C for 1 h prior to cutting 12-µm sections using a cryostat (Leica Biosystems Inc., Buffalo Grove, IL, USA). Sections were placed on positively charged slides, which were then stored at -20°C until they were shipped on dry ice to SUNY Upstate Medical University for immunohistochemical analysis. Slides were incubated on a 37°C heating block for 10 minutes. Sections were then permeabilized with 0.5% Triton^TM^ X-100, blocked with 7% goat serum (Gibco; Thermo Fisher Scientific), and incubated with primary antibodies followed by incubation with fluorescent secondary antibodies (**Table 2**); nuclei were counterstained with DAPI (Abcam). Slides were mounted with ProLong^TM^ Gold Antifade (Thermo Fisher Scientific), and fluorescent images were acquired with a Zeiss LSM780 confocal microscope (Zeiss, Germany). The image size was set to 1024 x 1024 pixels in x/y with a resolution of 3.85 µm per pixel. Individual z-stacks were captured with the z-step interval set to 0.5 µm. Fluorescence intensity of αSMA was measured in a standardized region of interest (i.e., SC inner wall and filtering TM region) using maximum intensity projections in FIJI software (NIH) with image background subtraction, followed by calculation of fold-change vs. control.

### Statistical analysis

Individual sample sizes are specified in each figure caption. Comparisons between groups were assessed by unpaired or paired t-tests, and one-way or two-way analysis of variance (ANOVA) with Tukey’s multiple comparisons *post hoc* tests, as appropriate. The significance level was set at p<0.05 or lower. GraphPad Prism software v10.0.2 (GraphPad Software, La Jolla, CA, USA) was used for all analyses.

## Results

### ECM stiffening induces SC cell YAP activity and cytoskeletal remodeling, which is reversed with matrix softening

The TM from glaucoma eyes is ∼1.5-20-fold stiffer compared to that from normal eyes (6-9). As such, SC inner wall cells in the diseased outflow tract are exposed to increased biomechanical stress from their underlying stiffened, ECM-rich substrate. We designed a hybrid ECM-alginate hydrogel that allows for dynamic changes of elastic modulus independent of ECM composition while maintaining cell-biomaterial interactions (**Fig. 1A**). ECM-alginate hydrogel stiffness was significantly increased by 3.3-fold with Ca^2+^-mediated crosslinking compared to controls, in line with reported differences in the glaucomatous TM tissue. Upon treatment with alginate lyase, hydrogel stiffness significantly decreased reaching soft baseline levels (**Fig. 1B**). As such, our hybrid ECM-alginate hydrogel facilitates dynamic bidirectional manipulation of matrix stiffness for interrogations of SC cellular responses.

**Fig. 1.**
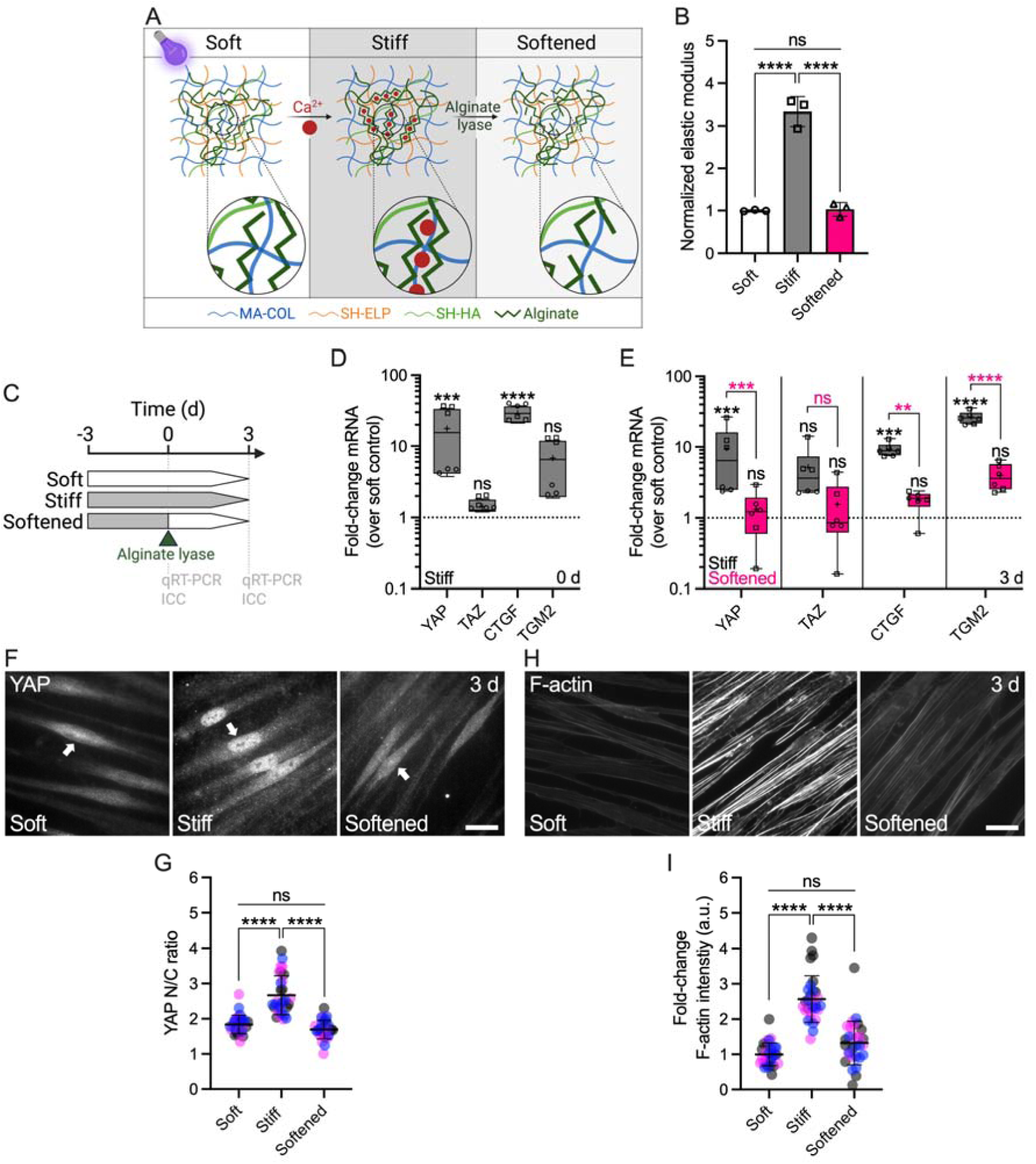
ECM stiffening induces YAP activity and F-actin remodeling in SC cells, which is reversed with matrix softening. (A) Schematic of calcium-mediated ECM-alginate hydrogel stiffening and alginate lyase-mediated matrix softening. (B) Normalized elastic modulus (to soft controls) by oscillatory rheology analysis (N = 3 experimental replicates per group). (C) Schematic showing the time course of *in situ* matrix softening with soft and stiff controls. (D) Normalized mRNA fold-changes (to soft controls) by qRT-PCR at 0 d and (E) 3 d (N = 6 experimental replicates per group from 2 HSC cell strains). (F) Representative fluorescence micrographs of YAP and (H) F-actin. Scale bars, 20 μm; arrows indicate YAP nuclear localization. (G) Analysis of YAP nuclear/cytoplasmic ratio and (I) F-actin fluorescence intensity (N = 30 images per group from 3 HSC cell strains with 3 experimental replicates per cell strain). Symbols with different shapes/colors represent different cell strains. The bars and error bars indicate mean ± SD. Box plots display median as horizontal line, interquartile range as boxes, and minimum/maximum range as whiskers; mean values are indicated by +. Significance was determined by one-way or two-way ANOVA using multiple comparisons tests (*p < 0.05; **p < 0.01; ****p < 0.0001; ns = non-significant difference).

To investigate the influence of substrate stiffness on YAP mechanosignaling, SC cells were (i) cultured on stiff hydrogels for 3 d before (ii) dynamically decreasing elastic modulus at 0 d using alginate lyase, (iii) followed by further culture on the softened matrix for 3 d; unmodified soft and stiff hydrogels served as controls (**Fig. 1C**). First, we observed that SC cells on stiff hydrogels at day 0 (i.e., 3-day exposure) had significantly increased mRNA levels of YAP and the known downstream effector connective tissue growth factor (CTGF) compared to soft controls, whereas the YAP paralog transcriptional coactivator with PDZ-binding motif (TAZ) and the putative effector transglutaminase 2 (TGM2) were not significantly altered (**Fig. 1D**). Immunostaining showed significantly increased YAP and TAZ nuclear translocation, the principal mechanism regulating their function, on stiff hydrogels compared to soft controls at day 0 (**Suppl. Fig. 1**).

Next, we tested whether stiff-induced YAP signaling could be reversed by matrix softening. SC cells on stiff hydrogels at day 3 (i.e., 6-day exposure) showed significantly increased YAP, CTGF, and TGM2 mRNA levels compared to soft controls, with TAZ expression not being significantly different. These alterations were potently reversed upon matrix softening; all transcript levels in the softened group were significantly decreased compared to SC cells on stiff hydrogels, matching soft controls (**Fig. 1E**). Immunostaining showed significantly increased YAP nuclear localization (**Fig. 1F,G**) and filamentous (F)-actin stress fibers (**Fig. 1G,H**) in SC cells on stiff hydrogels compared to soft controls. Matrix softening significantly decreased nuclear YAP levels and F-actin stress fibers - indistinguishable from soft controls - compared to SC cells on stiff hydrogels (**Fig. 1F-H**). Similarly, the stiff-induced expression of TGM2 was completely abolished with hydrogel softening (**Suppl. Fig. 2**).

Together, these data suggest that pathologic YAP activation and cytoskeletal reorganization in SC cells induced by short-term ECM stiffening is fully reversible by matrix softening.

### Reversal of stiff-induced SC cell YAP activity and cytoskeletal remodeling with matrix softening exhibits distinct temporal sequence

A recent study demonstrated rapid YAP nuclear-to-cytoplasmic redistribution in human mesenchymal stem cells as early as 0.5 h after ECM softening (44). Therefore, we next investigated the time-dependent reversal of stiff-induced (i.e., for 3 d) YAP nuclear localization and F-actin reorganization in SC cells 3 h, 1 d, 3 d, and 5 d after matrix softening (**Fig. 2A**). Consistent with earlier observations, SC cells on stiff hydrogels showed significantly increased nuclear YAP and F-actin stress fibers compared to soft controls at day 0. The YAP nuclear-to-cytoplasmic ratio was reversed to soft baseline levels as early as 3 h after matrix softening and then stabilized (**Fig. 2B,C**). In contrast, F-actin stress fiber intensity decreased more slowly; a significant difference compared to SC cells on stiff hydrogels was observed only at 24 h post-softening (**Fig. 2D,E**). Of note, nuclear YAP and F-actin stress fiber levels remained unchanged over time in SC cells on soft (i.e., low) or stiff hydrogels (i.e., high), respectively (**Suppl. Fig. 3**).

**Fig. 2.**
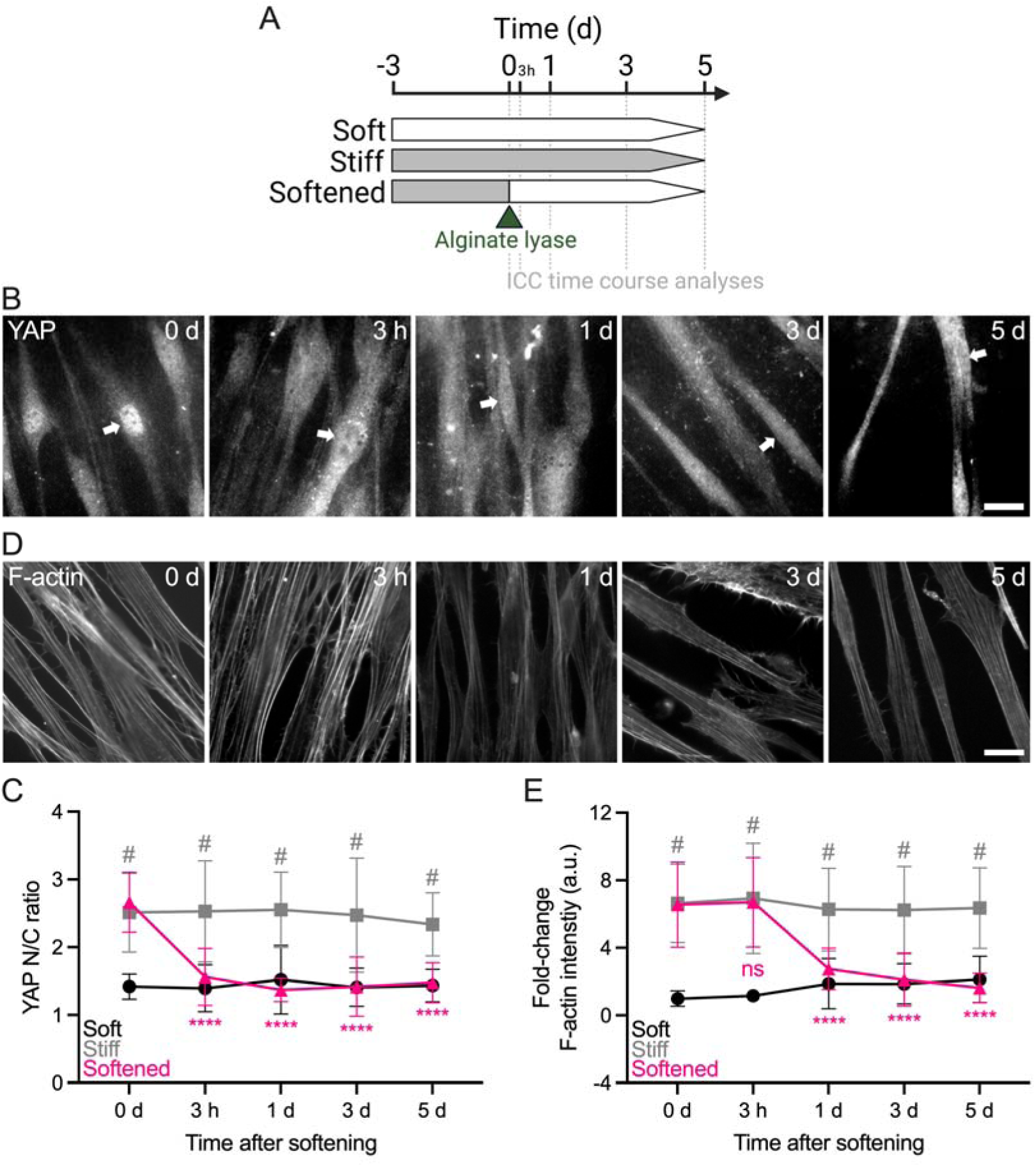
Reversal of stiff-induced YAP activity and F-actin remodeling in SC cells with matrix softening exhibits distinct temporal sequence. (A) Schematic showing the time course of stiff ECM-alginate hydrogel softening using alginate lyase. (B) Representative fluorescence micrographs of YAP and (D) F-actin over time. Scale bars, 20 μm; arrows indicate YAP nuclear localization. (C) Analysis of YAP nuclear/cytoplasmic ratio and (E) F-actin fluorescence intensity (N = 20 images per group from 2 HSC cell strains with 3 experimental replicates per cell strain). The bars and error bars indicate mean ± SD. Significance was determined by two-way ANOVA using multiple comparisons tests (****p < 0.0001 versus stiff at 0 d; ^#^p < 0.0001 versus soft per time point; ns = non-significant difference).

These data suggest that hyperactivation of YAP in SC cells stimulated by short-term ECM stiffening is rapidly rescued by matrix softening, whereas actin cytoskeletal adaptions occur in a slightly more delayed manner.

### Disruption of YAP signaling attenuates stiff-induced SC cell dysfunction

YAP lacks DNA-binding domains and binds primarily to TEAD transcription factors to drive stiff-responsive gene expression (15, 16). To investigate the effects of inhibiting YAP-TEAD interaction on SC cell pathobiology, cells were cultured on stiff hydrogels for 3 d in presence or absence of verteporfin (**Fig. 3A**). Verteporfin induces sequestration of YAP in the cytoplasm via increasing levels of the chaperone 14-3-3σ (45), thereby blocking transcriptional activation in ocular (21, 27) and non-ocular cells (46). After 3 d of verteporfin treatment, SC cells exhibited significantly decreased mRNA levels of YAP and TAZ compared to vehicle controls (**Fig. 3B**). Immunostaining confirmed significantly reduced YAP nuclear localization in SC cells on stiff hydrogels exposed to verteporfin compared to controls – validating its inhibitory mode of action (45) – concurrent with significantly decreased F-actin stress fibers (**Suppl. Fig. 4**). Similar observations were made for the downstream effectors CTGF and TGM2, on both mRNA and protein levels (**Fig. 3B-D**). YAP inhibition with verteporfin nearly abolished fibronectin (FN) mRNA expression, and significantly decreased protein deposition (**Fig. 3B,E,F**). Lastly, after 3 d of verteporfin treatment, SC cells showed significantly decreased alpha smooth muscle actin (αSMA) transcript and protein levels as well as phospho-myosin light chain (p-MLC) expression compared to vehicle controls (**Fig. 3B,G-J**).

**Fig. 3.**
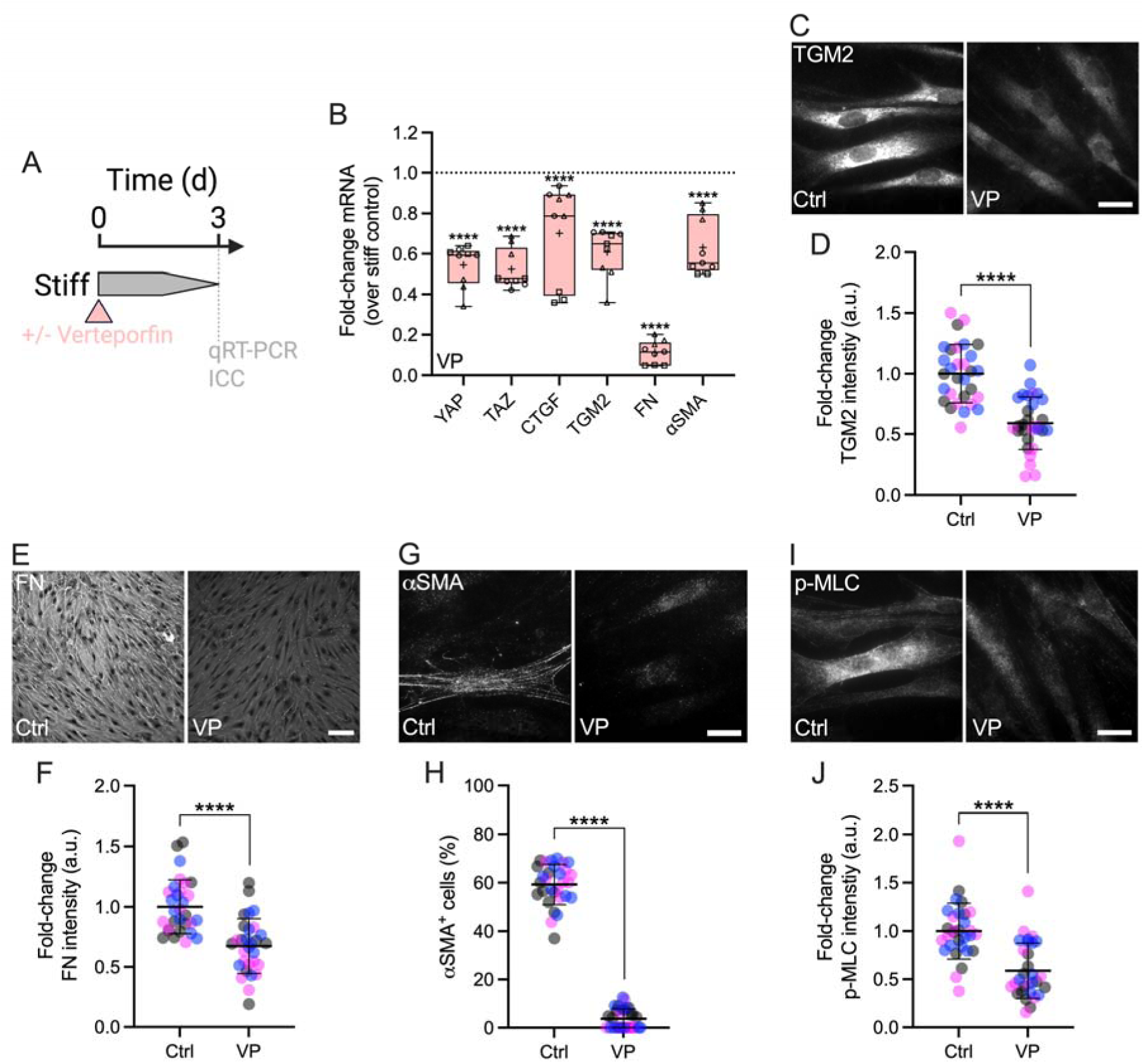
Interference with YAP-TEAD interactions attenuates stiff-induced dysfunction in SC cells. (A) Schematic showing the time course of pharmacologic YAP-TEAD inhibition with verteporfin (VP) in SC cells on stiff ECM-alginate hydrogels. (B) Normalized mRNA fold-changes (to stiff controls) by qRT-PCR at 3 d (N = 9 experimental replicates per group from 3 HSC cell strains). (C) Representative fluorescence micrographs of TGM2, (E) FN, (G) αSMA, and (H) p-MLC. Scale bars, 20 μm. (D) Analysis of TGM2, (F) FN, (H) number of αSMA^+^ cells, and (I) p-MLC fluorescence intensity (N = 30 images per group from 3 HSC cell strains with 3 experimental replicates per cell strain). Symbols with different shapes/colors represent different cell strains. The bars and error bars indicate mean ± SD. Box plots display median as horizontal line, interquartile range as boxes, and minimum/maximum range as whiskers; mean values are indicated by +. Significance was determined by two-way ANOVA using multiple comparisons tests and unpaired t-tests (****p < 0.0001).

To validate our findings using pharmacologic YAP inhibition, we knocked down YAP and TAZ using combined siYAP/TAZ in SC cells on stiff hydrogels (**Fig. 4A**), shown to be more effective in depleting their protein levels compared to the respective single siRNA treatments (**Suppl. Fig. 5A-C**). siYAP/TAZ transfection significantly reduced both YAP and TAZ mRNA compared to non-targeting siRNA controls, with TAZ transcript levels being lowered to a greater extent compared to YAP (**Fig. 4B**). Immunoblot analyses showed significantly decreased YAP and TAZ protein expression – again with TAZ being more affected than YAP – in agreement with the mRNA findings (**Fig. 4C-E**). Furthermore, immunostaining demonstrated significantly reduced YAP nuclear localization in SC cells on stiff hydrogels treated with siYAP/TAZ compared to controls (**Fig. 4F,G**), concurrent with significantly decreased F-actin stress fibers (**Suppl. Fig. 5D,E**). siYAP/TAZ treatment significantly reduced mRNA and protein levels of the downstream effectors CTGF and TGM2 compared to control siRNA, consistent with the verteporfin data (**Fig. 4B,H,I**). Similarly, siYAP/TAZ-treated SC cells showed significantly decreased FN mRNA expression and protein deposition (**Fig. 4B; Suppl. Fig. 5F,G**). Significantly decreased αSMA transcript and protein levels, as well as p-MLC expression were observed compared to vehicle controls (**Fig. 4B,J,K; Suppl. Fig. 5H,I**). Of note, the mRNA expression of canonical Hippo pathway kinases LATS1 and LATS2 in SC cells, together with the YAP chaperone 14-3-3σ was not affected by siYAP/TAZ knockdown (**Fig. 4B)**.

**Fig. 4.**
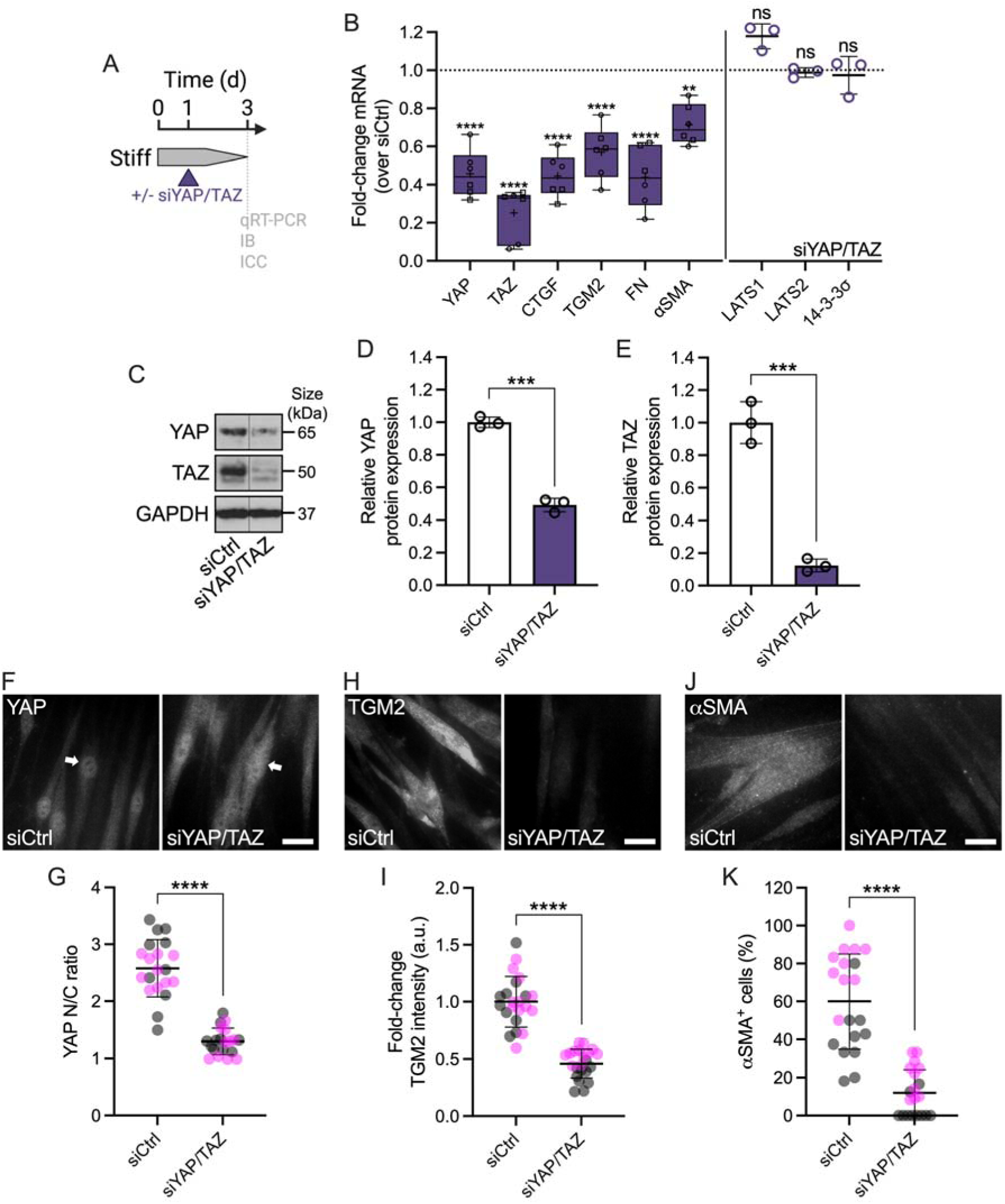
Genetic YAP (and TAZ) depletion attenuates stiff-induced dysfunction in SC cells. (A) Schematic showing the time course of siRNA-mediated YAP/TAZ depletion in SC cells on stiff ECM-alginate hydrogels. (B) Normalized mRNA fold-changes (to control siRNA) by qRT-PCR at 3 d (N = 6 experimental replicates per group from 2 HSC cell strains). (C) Representative immunoblot of YAP and TAZ with GAPDH serving as loading control. (D) Analysis of relative YAP and (E) TAZ protein levels (normalized to GAPDH) (N = 3 experimental replicates per group from 1 HSC cell strain). (F) Representative fluorescence micrographs of YAP, (H) TGM2, and (J) αSMA. Scale bars, 20 μm; arrows indicate YAP nuclear localization. (G) Analysis of YAP nuclear/cytoplasmic ratio, (I) TGM2 fluorescence intensity, and (K) number of αSMA^+^ cells (N = 20 images per group from 2 HSC cell strains with 3 experimental replicates per cell strain). Symbols with different shapes/colors represent different cell strains. The bars and error bars indicate mean ± SD. Box plots display median as horizontal line, interquartile range as boxes, and minimum/maximum range as whiskers; mean values are indicated by +. Significance was determined by two-way ANOVA using multiple comparisons tests and unpaired t-tests (**p < 0.01; ***p < 0.001; ****p < 0.0001; ns = non-significant difference).

Together, these data suggest (i) that YAP is a central regulator of the SC cell pathologic response to ECM stiffening that involves altered cytoskeletal and ECM remodeling, and (ii) that pharmacologic or genetic disruption of YAP signaling potently attenuates glaucoma-like SC cell dysfunction induced by ECM stiffening.

### Pharmacologically targeting YAP signaling increases ex vivo outflow facility

We recently showed that verteporfin attenuates pathological contraction and stiffening of TM cell-encapsulated hydrogels (27), comparable to ROCK inhibitor treatment (32). Therefore, we hypothesized that pharmacologically targeting YAP signaling increases outflow facility through the conventional outflow pathway, potentially via tissue relaxation. To investigate the effects of inhibiting YAP-TEAD interaction on outflow function, anterior chambers of paired enucleated mouse eyes were cannulated, perfused for 20 minutes, and then exchanged with vehicle or verteporfin (**Fig. 5A**). Following the dual needle exchange regimen, perfusions were restarted; results showed that verteporfin treatment significantly increased *ex vivo* outflow facility by 38.3% compared to vehicle controls (**Fig. 5B**). No differences in gross morphology of the iridocorneal angle were observed between groups (**Fig. 5C; Suppl. Fig. 6**). However, immunostaining showed significantly decreased αSMA expression levels in the SC inner wall and underlying TM with verteporfin perfusion compared to vehicle controls (**Fig. 5D,E**).

**Fig. 5.**
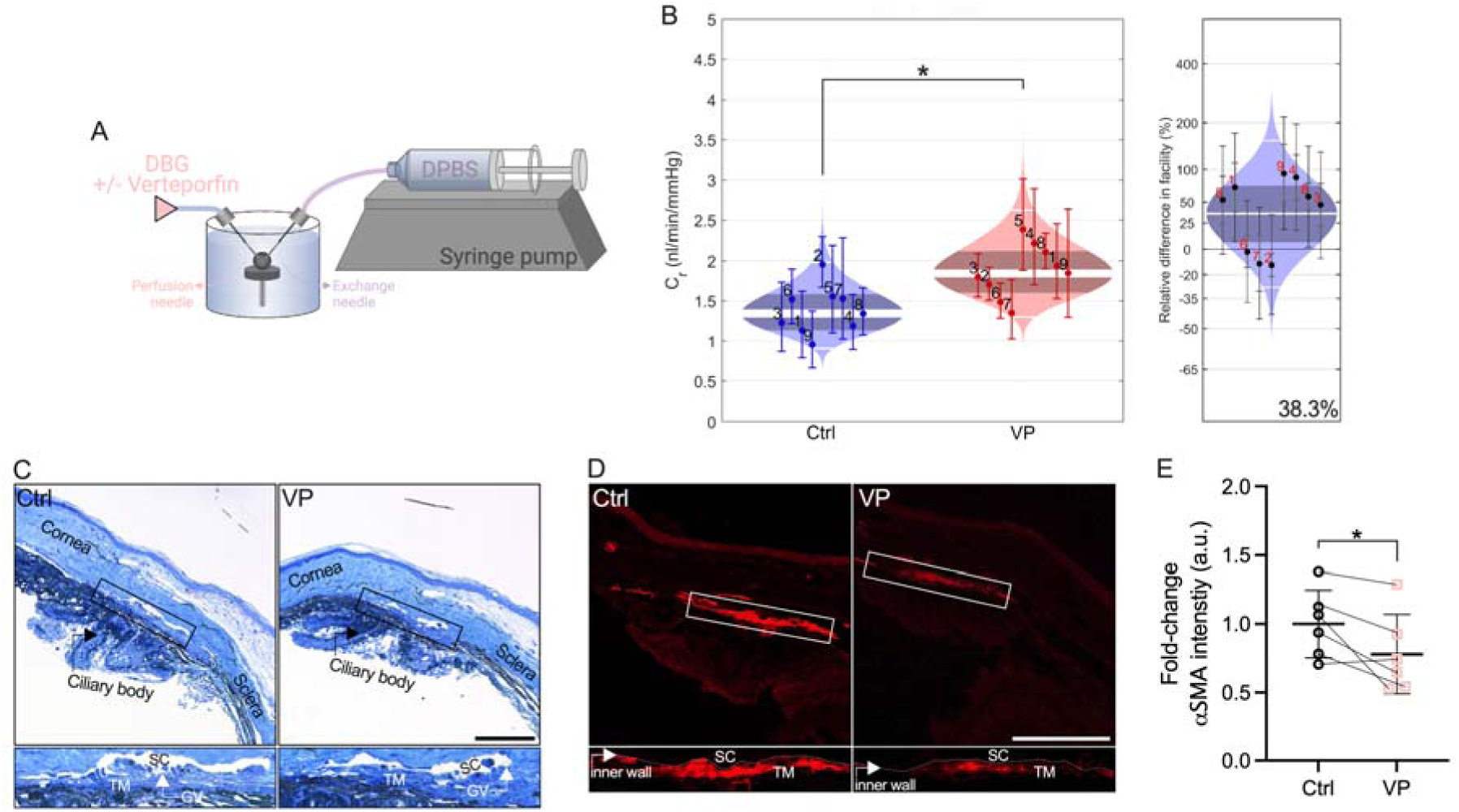
Pharmacologically targeting YAP signaling increases *ex vivo* outflow facility. (A) Schematic showing the core *iPerfusion* setup for dual-needle drug exchange. (B) Analysis of outflow facility including relative difference between groups (N = 9 eyes per group; paired). (C) Representative light micrographs of iridocorneal tissue and (D) representative fluorescence micrographs of αSMA. Scale bars, 100 μm; boxes show region of interest; GV = giant vacuole. (E) Paired analysis of αSMA fluorescence intensity (N = 6 images per group from 3 experimental replicates with two quadrants per sample). The bars and error bars indicate mean ± SD. Cello plots display predicted log-normal distribution as light shaded bands, the mean as central white line, the 95% CI on the mean as dark shaded bands, and ± two SDs around the mean as outer white lines. Facility for each eye with 95% CI is shown by individual data points and error bars. Significance was determined by paired t-tests (*p < 0.05).

Together, these data suggest that pharmacologic disruption of YAP signaling with a clinically-used small molecule is a potential strategy to increase outflow function via targeting cytoskeletal remodeling.

## Discussion

Pathologically altered biomechanical properties of the SC inner wall microenvironment, including stiffening of the underlying TM, were recently validated as the culprit for increased outflow resistance in ocular hypertensive glaucoma cases (9, 10). However, the involvement of specific mechanotransduction pathways in these disease processes is largely unclear. Here, we demonstrate that YAP is a central regulator of glaucoma-like SC cell dysfunction in response to ECM stiffening, and that targeted disruption of YAP mechanosignaling attenuates SC cell pathobiology and enhances outflow facility.

Previous studies have established a critical role of YAP mechanotransduction in normal eye development and disease, including glaucoma (47-49). In this context, the anatomical substrate of the SC inner wall – the TM – has been studied extensively in the last decade (19-26). Our group confirmed and extended prior findings of YAP mechanosignaling in TM cell dysfunction with emphasis on studying cellular behaviors in a tissue-like 3D ECM microenvironment (27, 28). Despite their close functional interdependence, the SC cell mechanobiology landscape in health and disease is considerably less well understood. We recently demonstrated, for the first time, that SC cells exhibit a remarkably similar YAP “signature” compared to TM cells when challenged with transforming growth factor beta2 (31). In the present study, we sought to elucidate SC cell YAP modulation in response to alterations in ECM stiffness.

The TM in glaucomatous eyes is markedly stiffer compared to that from normal eyes (6-9), thereby exerting increased biomechanical stress on SC inner wall cells *in situ*. To model this, we modified our bioengineered modular hydrogel system (32) and developed a novel hybrid ECM-alginate version. This composite hydrogel facilitates on-demand and reversible control over elastic modulus during the culture of cells to investigate how YAP mechanotransduction in SC cells is regulated by dynamic matrix stiffness changes. To tune hydrogel stiffness independent of ECM composition and without compromising cell-biomaterial interactions, additional bioinert polymers can be incorporated. They interlace with the ECM proteins and form a so-called interpenetrating network (50, 51). Alginate, derived from seaweed, is a widely-used biocompatible polymer with a gentle mode of crosslinking by soluble calcium or other divalent cations such as strontium. It presents no intrinsic cell-binding domains and therefore acts as a bioinert crosslinker (52, 53). For instance, collagen-alginate (54-58), fibrin-alginate (59), and polyethylene glycol-alginate (60) hydrogels have been described to investigate epigenetic reprogramming of tumor cells, cell migration and spreading, as well as mesenchymal stem cell differentiation. In addition, hybrid ECM-alginate hydrogels proved their utility to model tissue properties in several ocular and neural diseases (61-64). A key feature of alginate from a materials perspective is its bidirectional responsiveness to crosslinking (= stiffening) and degradation (= softening). Our data showed that Ca^2+^-stiffened hydrogel elastic modulus was fully reversed to soft baseline levels upon alginate degradation in a cytocompatible manner using alginate lyase (65).

Increased substrate stiffness is a potent driver of YAP nuclear localization; in the nucleus, YAP binds primarily to TEAD transcription factors to drive stiff-responsive gene expression (15, 16). Our data showed that ECM stiffening increased YAP transcript in SC cells, while TAZ expression was unaffected. A similar pattern was observed for mRNA levels of CTGF (= up) and TGM2 (= unchanged) – two prominent effectors of active YAP/TAZ signaling that have been implicated in outflow dysfunction in glaucoma (66, 67). Although we found their mRNAs to be differentially expressed, the stiff-induced YAP and TAZ nuclear translocation response – the principal mechanism regulating their activity – was nearly identical. This was in agreement with their acknowledged functional redundancy (68) and suggests that the two paralogs may play overall analogous roles in the context of SC cell mechanotransduction, as they do in TM cells (27).

Having established that ECM stiffening stimulates aberrant YAP mechanosignaling, we next asked whether these induced pathological alterations could be reversed by matrix softening. Our data showed that stiff-induced YAP activation, i.e., nuclear translocation and transcriptional activation (as evidenced by TGM2 expression), is fully reversible with matrix softening. YAP mechanotransduction requires actomyosin cytoskeletal integrity (69, 70). We found that cytoskeletal disorganization in SC cells induced by short-term ECM stiffening was potently reversed with alginate lyase treatment; contractile F-actin stress fibers, whose role is to counteract the mechanical forces acting upon them, completely disassembled reaching soft baseline controls. Next, we sought to investigate the dynamics of the YAP/F-actin adaption processes after the hydrogel modulus was softened. Our data showed that YAP quickly translocated from the nucleus to the cytoplasm as early as 3 h after ECM softening and then stabilized. This rapid adaptation of YAP activity to matrix softening is consistent with previous studies that reported similar sensitivity of YAP signaling to mechanical cues in mesenchymal stem cells (44). We observed an interesting phenomenon in which F-actin levels decreased at a slower rate compared to YAP nuclear localization, which took 24 h to reach soft baseline levels. The polymerization and depolymerization of actin filaments is a dynamic process controlled by a variety of regulatory proteins (71). Among them, Rho GTPases are essential mediators connecting mechanical stimuli and actin-dependent YAP regulation (17). Rho stimulates the assembly of contractile actin stress fibers – composed of F-actin and non-muscle myosin II – by activating downstream effectors such as Rho-associated kinase and mDia1/2 (72). The apparent delay in F-actin stress fiber disassembly in SC cells upon matrix softening relative to YAP cytoplasmic redistribution could be explained by potentially slowed transition between filamentous (F-actin) and monomeric (G-actin) states under the control of nucleotide hydrolysis, or altered actin binding protein regulation in the serum-free experimental conditions (73).

Targeting ECM mechanics by preventing or reversing tissue stiffening is an emerging therapeutic approach with clear implications for treatment of ocular hypertension in glaucoma (74). The photosensitizer verteporfin has long been used clinically to treat age-related macular degeneration via photodynamic therapy using non-thermal laser light (75). In addition, verteporfin has demonstrated activity independent of light-activation; it potently inhibits YAP-TEAD interaction to block transcriptional activation (46) by sequestering YAP in the cytoplasm via increasing levels of the chaperone 14-3-3σ (45). We recently demonstrated that verteporfin blocked pathological YAP activation and potently mitigated transforming growth factor beta2-induced ECM remodeling, contraction, and stiffening in both TM and SC cells (27, 31). Our data here showed that pharmacologic (and confirmatory genetic) disruption of YAP signaling with verteporfin attenuates glaucoma-like SC cell dysfunction induced by ECM stiffening. We found this to occur on both the mRNA and protein levels, supportive of functional YAP interference. Specifically, fibrotic-like actomyosin cytoskeletal remodeling and dysregulated ECM deposition/crosslinking – akin to endothelial-to-mesenchymal transition (76) – was completely attenuated with verteporfin.

With compelling *in vitro* data in hand, we then tested whether targeted disruption of YAP mechanosignaling using verteporfin increases *ex vivo* outflow facility. Mouse eyes are an ideal model for investigating outflow function owing to the anatomical relationship between the TM and inner wall of SC, and demonstrated bidirectional responses to outflow-altering drugs comparable to human eyes (77-79). Using the *iPerfusion* system (40) in drug exchange regime, our data showed that verteporfin perfusion increased outflow facility by almost 40% over controls. The iridocorneal angles in drug-treated eyes appeared grossly normal, in fact indistinguishable from controls. By contrast, we found decreased expression of αSMA (the actin isoform typical of vascular smooth muscle cells) in the immediate vicinity of the SC inner wall, especially in cells throughout the filtering TM region, consistent with previous reports (80, 81). These combined data suggest that the increase in outflow facility following inhibition of YAP-TEAD interaction with verteporfin is functionally mediated via targeting actin cytoskeleton dynamics. In support of this notion, the FDA-approved Rho-associated kinase inhibitor netarsudil increases outflow via cell/tissue relaxation, mediated in part by actin stress fiber disassembly (82, 83). Future studies are aimed at perfusing verteporfin in hypertensive eyes *ex vivo* for further validation of its use in “mechanomedicine”.

In conclusion, we demonstrate a pathologic role of aberrant YAP mechanosignaling in SC cell pathobiology and that pharmacologic disruption of YAP-TEAD interaction with the clinically-used small molecule verteporfin is a potential strategy to increase outflow facility for prevention or treatment of ocular hypertension in glaucoma.

## Acknowledgments

We thank Dr. Robert W. Weisenthal and the team at Specialty Surgery Center of Central New York for assistance with corneal rim specimens. We also thank Dr. Nasim Annabi at the University of California – Los Angeles for providing the KCTS-ELP, Dr. Alison Patteson at Syracuse University for rheometer access, and Dr. Audrey M. Bernstein and the Neuroscience Microscopy Core at SUNY Upstate Medical University for imaging support.

## Author contributions

H.L., M.K., R.A.K., A.S., K.K.P., I.S., M.L.De I., W.D.S., P.S.G., and S.H. designed all experiments, collected, analyzed, and interpreted the data. M.K., M.L.De I., and W.D.S. performed the *ex vivo* perfusion experiments and analyzed the data. R.A.K. and A.S. performed the histology and immunohistochemistry experiments and analyzed the data. H.L. and S.H. wrote the manuscript. All authors commented on and approved the final manuscript. W.D.S., P.S.G. and S.H. conceived and supervised the research.

## Data and materials availability

All data needed to evaluate the conclusions in the paper are present in the paper and/or the Supplementary Materials. Additional data related to this paper may be requested from the authors.

