## Supplementary Material for "Targeting YAP mechanosignaling to ameliorate stiffness-induced Schlemm’s canal cell pathobiology"

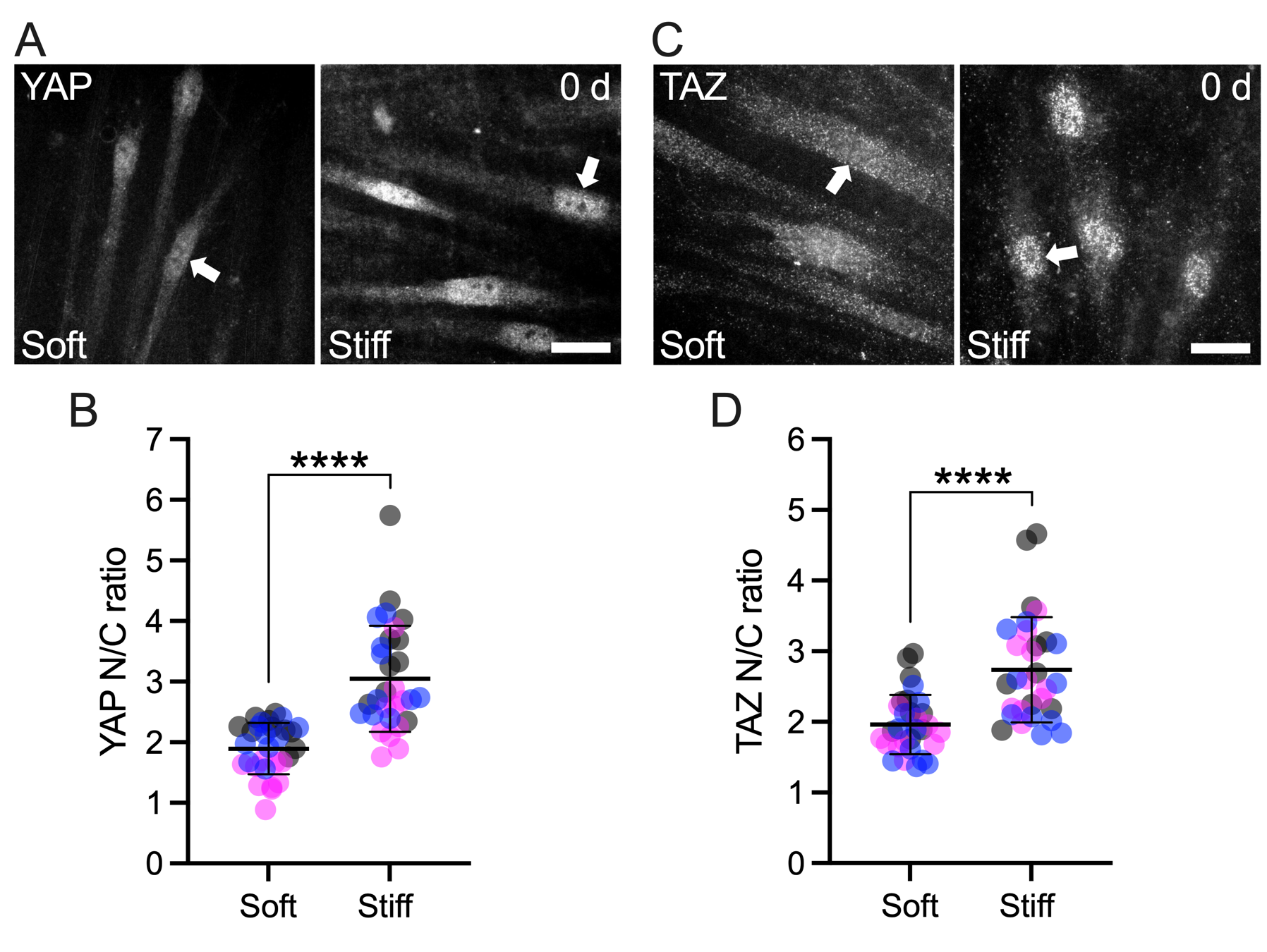

**Suppl. Fig. 1. ECM stiffening induces YAP and TAZ nuclear localization in SC cells.** (A) Representative fluorescence micrographs of YAP and (C) TAZ. Scale bars, 20 μm; arrows indicate YAP/TAZ nuclear localization. (B) Analysis of YAP and (D) TAZ nuclear/cytoplasmic ratios (N = 30 images per group from 3 HSC cell strains with 3 experimental replicates per cell strain). Symbols with different colors represent different cell strains. The bars and error bars indicate mean ± SD. Significance was determined by unpaired t-tests (****p < 0.0001).

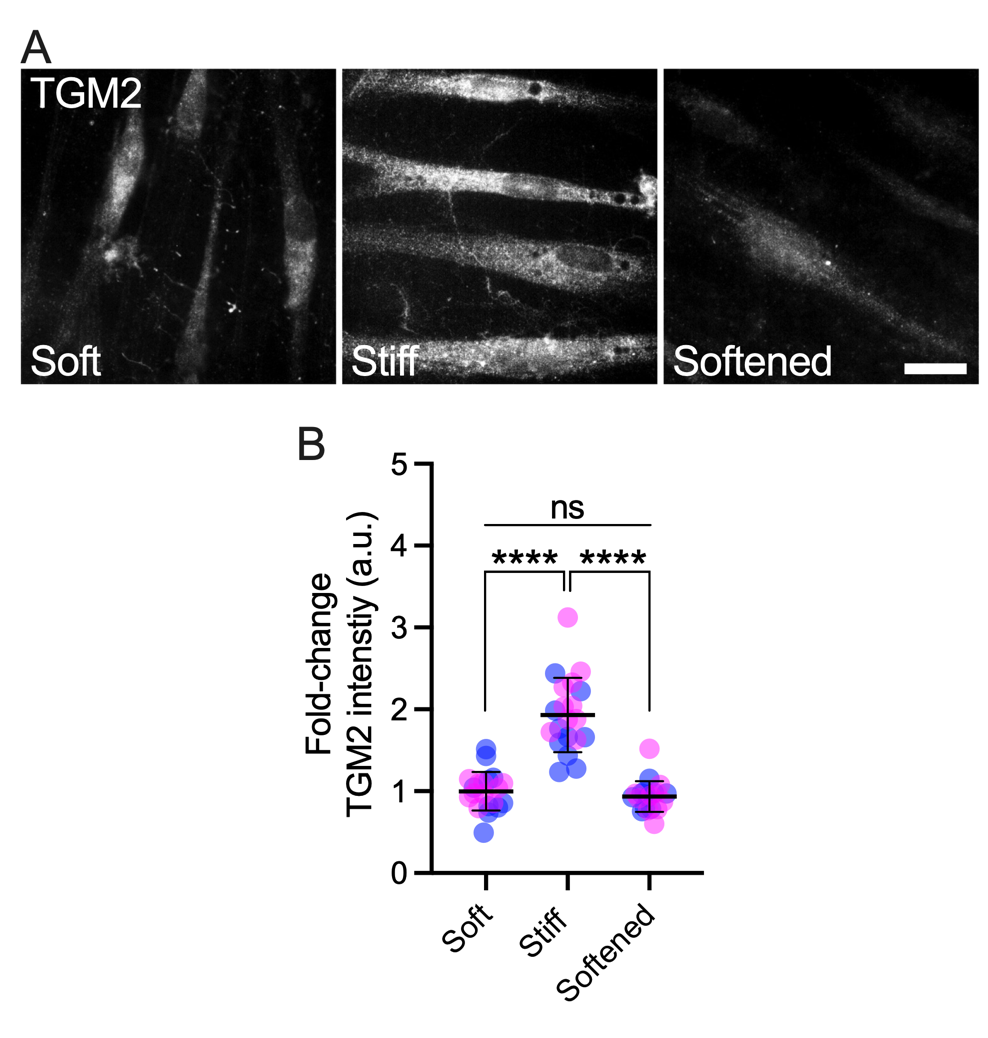

**Suppl. Fig. 2. ECM stiffening induces TGM2 levels in SC cells, which is reversed with matrix softening.** (A) Representative fluorescence micrographs of TGM2. Scale bar, 20 μm. (B) Analysis of TGM2 fluorescence intensity (N = 20 images per group from 2 HSC cell strains with 3 experimental replicates per cell strain). Symbols with different colors represent different cell strains. The bars and error bars indicate mean ± SD. Significance was determined by two-way ANOVA using multiple comparisons tests (****p < 0.0001; ns = non-significant difference).

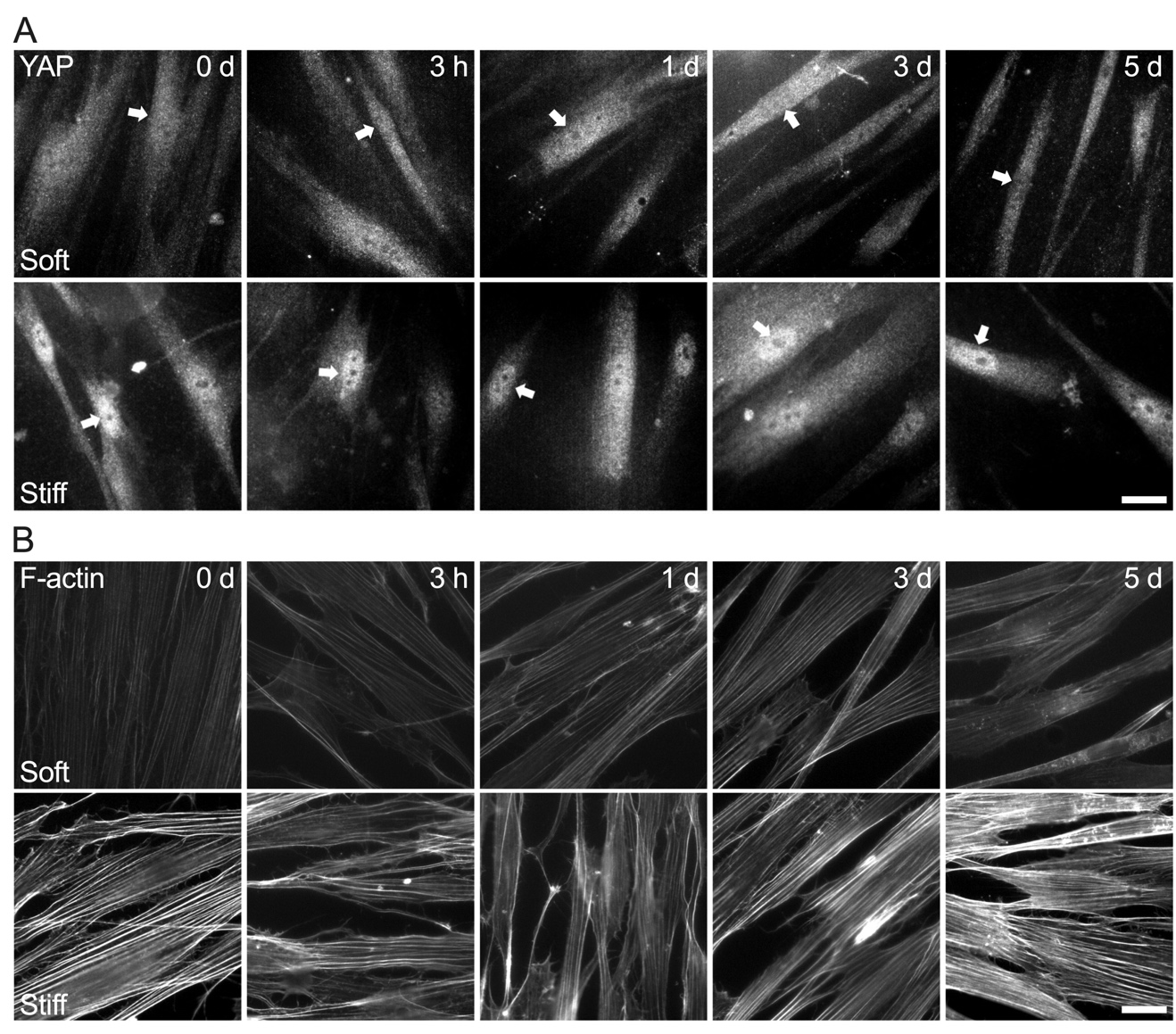

**Suppl. Fig. 3. YAP subcellular localization and F-actin organization in SC cells is dependent on hydrogel stiffness.**  (A) Representative fluorescence micrographs of YAP and (B) F-actin over time. Scale bars, 20 μm; arrows indicate YAP nuclear localization.

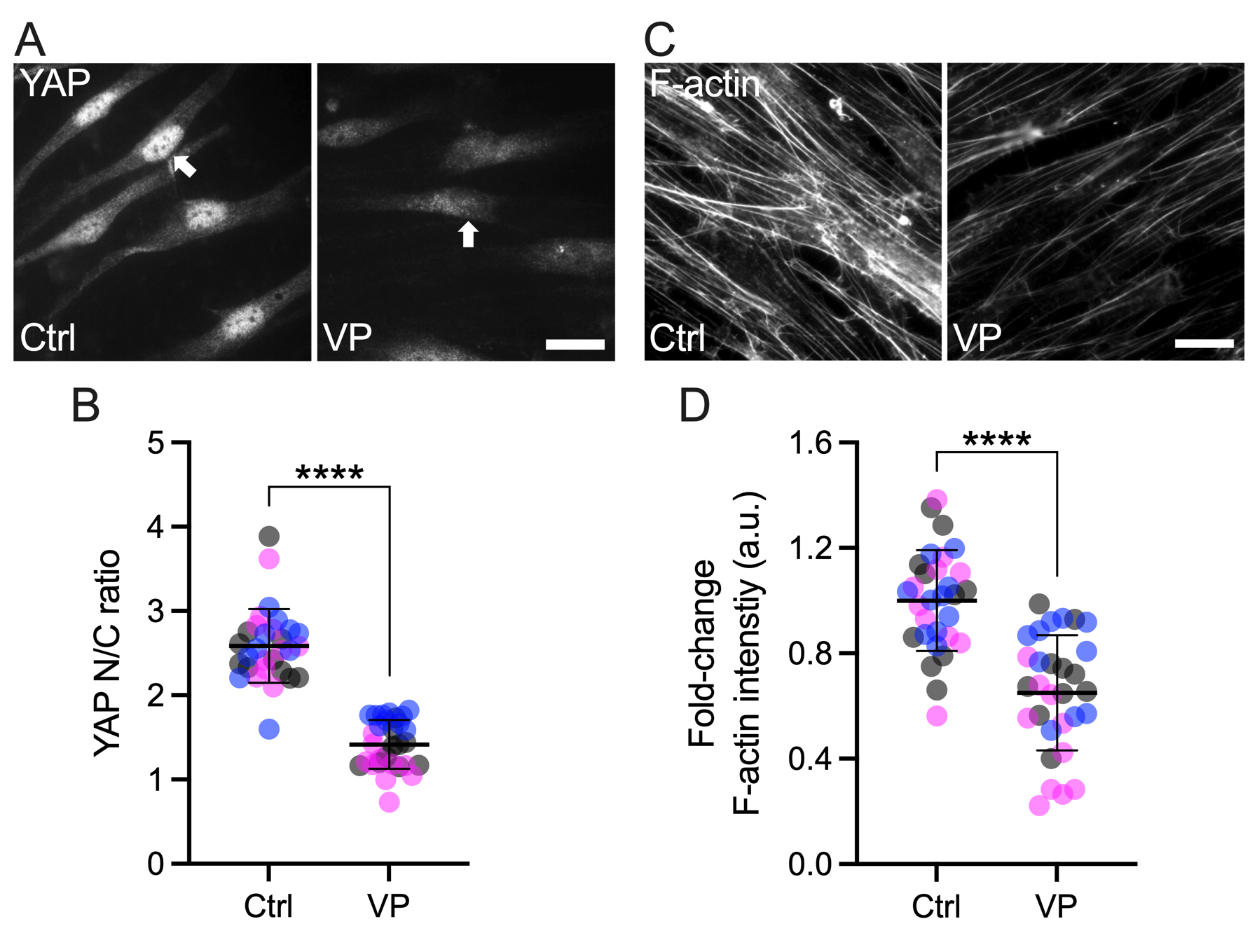

**Suppl. Fig. 4. Interference with YAP-TEAD interactions attenuates stiff-induced dysfunction in SC cells.** (A) Representative fluorescence micrographs of YAP and (C) F-actin. Scale bars, 20 μm; arrows indicate YAP nuclear localization. (B) Analysis of YAP nuclear/cytoplasmic ratios and (D) F-actin fluorescence intensity (N = 30 images per group from 3 HSC cell strains with 3 experimental replicates per cell strain). Symbols with different colors represent different cell strains. The bars and error bars indicate mean ± SD. Significance was determined by unpaired t-tests (****p < 0.0001).

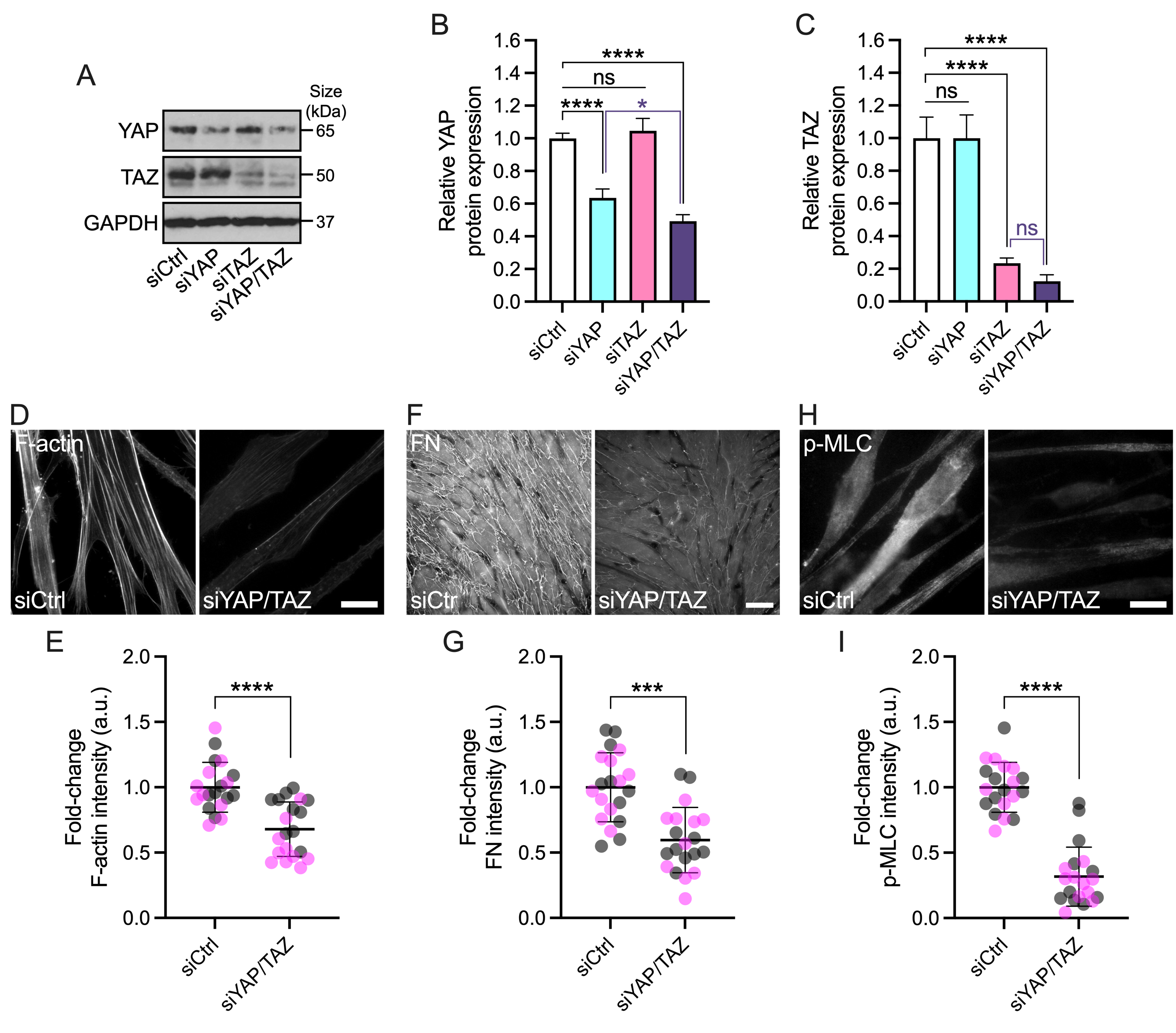

**Suppl. Fig. 5. Genetic YAP (and TAZ) depletion attenuates stiff-induced dysfunction in SC cells.** (A) Representative immunoblot of YAP and TAZ with GAPDH serving as loading control. (B) Analysis of relative YAP and (C) TAZ protein levels (normalized to GAPDH) (N = 3 experimental replicates per group from 1 HSC cell strain). (D) Representative fluorescence micrographs of F-actin, (F) FN, and (H) p-MLC. Scale bars, 20 μm. (E) Analysis of F-actin, (G) FN, and (I) p-MLC fluorescence intensity (N = 20 images per group from 2 HSC cell strains with 3 experimental replicates per cell strain). Symbols with different colors represent different cell strains. The bars and error bars indicate mean ± SD. Significance was determined by one-way ANOVA using multiple comparisons tests and unpaired t-tests (*p < 0.05; ***p < 0.001; ****p < 0.0001; ns = non-significant difference).

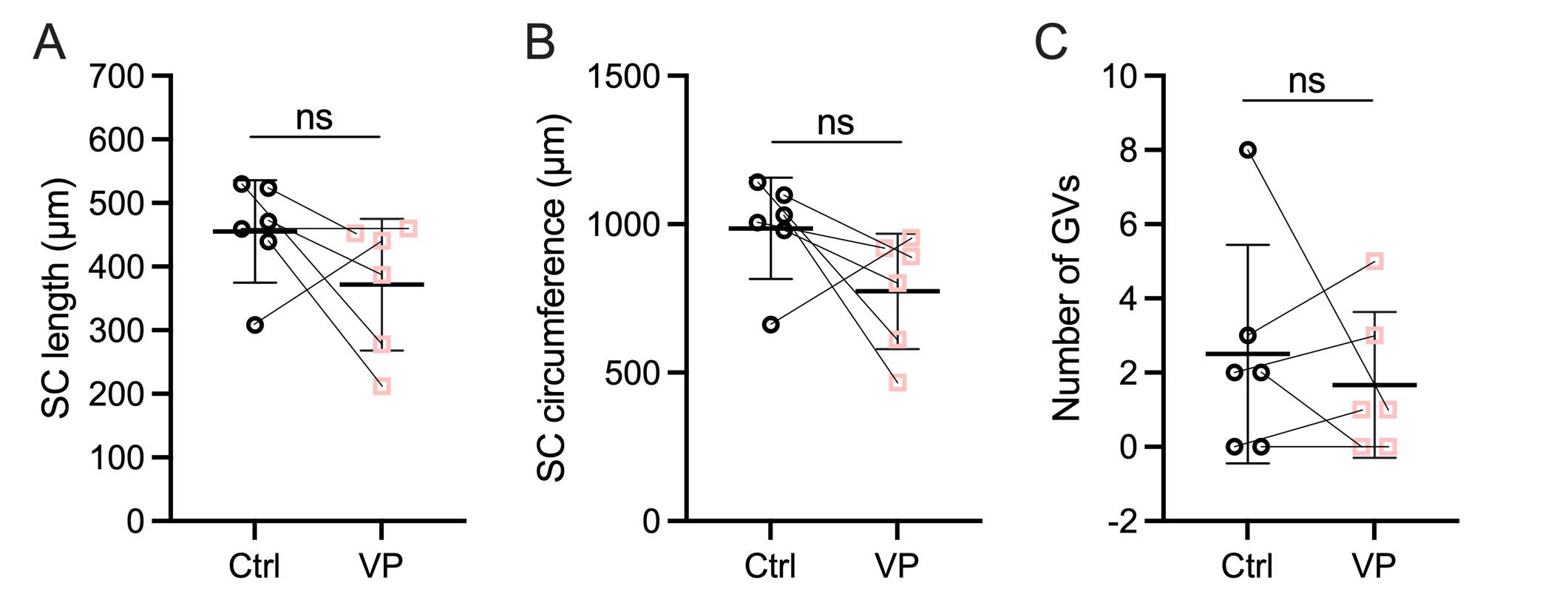

**Suppl. Fig. 6. Pharmacologically targeting YAP signaling does not alter gross SC morphology.** (A) Paired analysis of SC length, (B) SC circumference, and (C) number of giant vacuoles (GVs) (N = 6 images per group from 3 experimental replicates with two quadrants per sample). The bars and error bars indicate mean ± SD. Significance was determined by paired t-tests (ns = non-significant difference).
